# LRP-1 Commands Two Parallel Signalling Mechanisms to Support Constitutive Tumour Cell Motility in the Absence of Blood Supply

**DOI:** 10.1101/2021.09.10.459861

**Authors:** Cheng Chang, Xin Tang, Daniel Mosallaei, Mei Chen, David T. Woodley, Axel H. Schönthal, Wei Li

**Author notes:** These authors contributed equally to this work (Cheng Chang, Xin Tang).

## Abstract

Tumour cells often face the stress of ischemic (nutrient paucity and hypoxia) environment and must act self-sufficient to migrate toward the nearest blood supply or die. The mechanism that supports the constitutive motility of tumour cells under stress is poorly understood. We and others have previously shown that the low-density lipoprotein receptor-related protein 1 (LRP-1) plays a critical role in tumour cell migration and invasion *in vitro* and tumour formation in mice. Herein we show that depletion of LRP-1 completely abolishes the self-supported and serum-independent tumour cell motility. More intriguingly, we demonstrate that LRP-1 commands the full tumour cell motility by connecting with two independent cell surface signalling pathways. First, LRP-1 mediates secreted Hsp90α signalling via the “Hsp90α > LRP-1 receptor autocrine loop” for a half of tumour cell motility. Second, LRP1 stabilizes constitutively activated EGFR signalling that contributes the other half of tumour cell motility. Only combined inhibitions of the secreted Hsp90α autocrine and the EGFR signalling reproduces the effect of LRP1 down-regulation on constitutive tumour cell motility. This study reveals a novel mechanism of how tumour cells migrate in the absence of blood support.

## Introduction

Low density lipoprotein receptor-related protein 1 (LRP-1), also known as alpha-2-macroglobulin receptor (A2MR), apolipoprotein E receptor (APOER) or cluster of differentiation 91 (CD91), is a member of the LDL receptor superfamily found in the plasma membrane of cells that includes at least 11 structurally related members (LDL receptor, LRP1/TβRV, LRP-1b, LRP2/Megalin, VLDL receptor, LRP-3, LRP4/MEGF7, LRP5, LRP6, ApoE receptor 2/LRP8, LRP-9) (Hertz and Strickland, 2001). Among them, LRP-1 is the only member widely expressed in most cell types. Following transcription and translation, the encoded large preproprotein of LRP-1 is proteolytically processed to generate a 515-kDa extracellular and an 85-kDa intracellular subunits that are non-covalently bound to form the mature LRP-1 receptor at cell surface. LRP-1 is functionally diverse as historically a scavenger receptor for endocytosis and as more recently a *bona fide* receptor to bind a wide range of structurally and functionally diverse extracellular ligands and transduce their signals to intracellular signal pathways, most noticeably the Akt, NF-κB and Erk1/2 pathways. LRP-1 has been shown to play a role in regulation of tissue inflammatory reaction, tissue remodelling and cell survival during tissue injury (Shinohara et al., 2017). LRP-1 gene knockout led to embryonic lethality in mice (Herz et., 1992).

Previous studies indicated that LRP-1 plays critical roles in tumour cell migration and invasion *in vitro*, albeit via a wide variety of possible mechanisms (Willnow et al., 1999; Lillis et al., 2005; Xing et al., 2016). Bu’s group showed that miR-205 down-regulates the expression of LRP-1 by targeting sequences in the 3’UTR of LRP1 mRNA and caused decreased migration of both U87 and SK-LU-1 tumour cells (Song et al., 2009a). Furthermore, this group reported that LRP-1 is required for glioblastoma cell migration and invasion *via* inducing expression of the matrix metalloproteinases 2 and 9 *in vitro* (Song et al., 2009b). We have shown that downregulation of LRP-1 blocked MDA-MB-231 cell migration *in vitro* (Zou et al. 2016) and tumour formation in mice (Sahu et al., 2012). Berquand et al reported that LRP-1 plays a crucial role in MDA-MB-231 cell migration by modulating the membrane extension dynamic (Berquand et al., 2019). Apert-Collin and colleagues showed that LRP-1 silencing leads to reorganizations of the actin-cytoskeleton by inhibiting FAK activation, activating RhoA activity and MLC-2 phosphorylation, thus preventing cell migration in a 3-D configuration assay (Apert-Collin et al., 2017). However, a major complexity from these studies was the failure to define nature of “tumour cells’ constitutive motility” that plays a <u>direct</u> role in tumour invasion and metastasis *in vivo*, i. e. the self-supported motility under hypoxia and nutrient paucity due to lack of support from the nearest blood vessels. In the current study, we have investigated the molecular mechanism that provides support for the constitutive and self-supported motility of the triple negative breast cancer cells, MDA-MB-231, one of the most used tumour cell models *in vitro* and *in vivo* (appeared in the titles of 17,061 publications by August 24, 2021). Findings of this study could provide new insights into design of therapeutic approaches to block invasion and metastasis of certain human breast cancers.

## Results

### Self-supported breast cancer cell motility versus serum-dependent non-cancer breast epithelial cell motility

Constitutive motility even in the absence of blood supply is a critical ability of tumour cells to achieve continued invasion and metastasis (Oudin and Weaver, 2016). To investigate the mechanism of this important cancer cell property, we first define the so-called “self-supported and constitutive motility” of tumour cells by comparing the highly malignant human breast cancer cell line, MDA-MB-231 (https://www.atcc.org/products/htb-26), with the non-tumorigenic human epithelial cell line, HBL-100 (https://lincs.hms.harvard.edu/db/cells/51117/). We utilized the colloidal gold migration assay, arguably the most sensitive and accurate cell migration assay that detects and quantitates single cell motility under any given condition by a darkfield microscope with assistance from a computer software, giving rise to a statistical sum of the so-called “migration Index” (%) of (Materials and Methods) (Li et al, 2004; Cheng et al., 2008). As shown in **Figure 1**, each migration track (black area) represents the migrated trail of a single cell under indicated time and environmental conditions. For example, in the absence of serum factor support, the non-cancer HBL-100 cells were entirely unable to migrate (panel b, indicated by white dotted circle). Under the full support of serum factors, the non-cancer cells migrated to create large tracks (panels a). Similarly, under the full support of serum factors, MDA-MB-231 cells also migrated to create large tracks (panel c), as expected. However, the motility of MDA-MB-231 cells was largely self-supported, since it was only slightly reduced by serum removal (emphasized by red dotted circle). In the presence of hypoxia (1% O_2_), the tumour cell migration remained largely unchanged (panels e and f). Computer-assisted quantitation of randomly selected 15 independent images of the cell migration under indicated conditions is shown in **Figure 1B**. These results indicated that MDA-MB-231 cells have acquired a self-supported mechanism to maintain their motility, due to cut off of the serum support.

**Figure 1.**
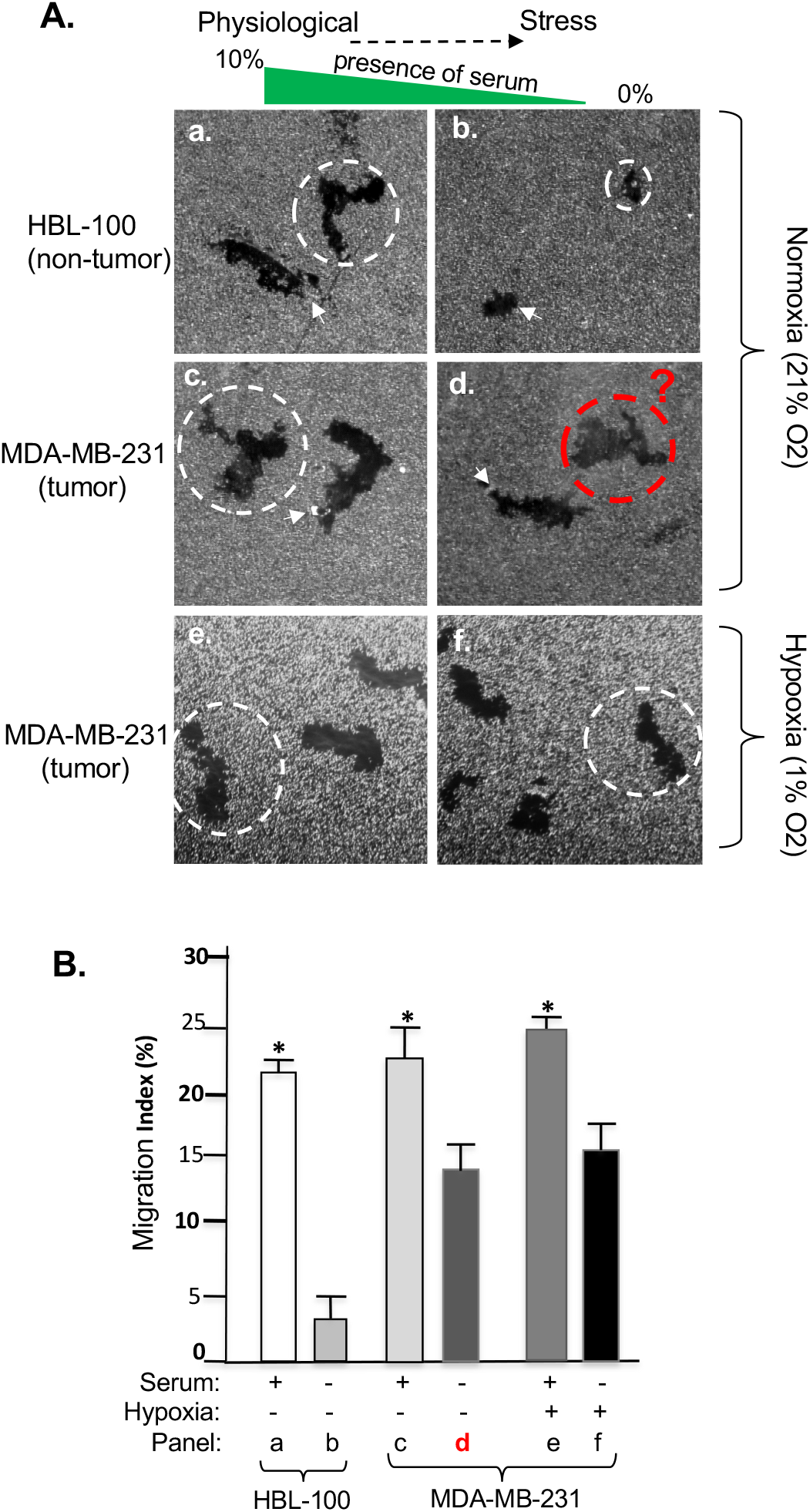
Self-supported constitutive motility of tumour, but not normal, cells. MDA-MB-231 and HBL-100 cells, cultured for three days in complete medium with 10% FBS to sub-confluence, were serum starved for 24 hours. Cells were lifted by trypsin, counted and subjected to colloidal gold migration assay in medium with or without 10% FBS or hypoxia (1% O2). A) Images of the representative cell migration tracks, whose average size was indicated by dotted circles. B) Computer-assisted quantitation of 15 randomly selected cell migration images for each experimental condition under a microscope, as Migration Index (%) (Methods). These results were confirmed by five independent experiments with statistical analyses, p < 0.05.

### LRP-1 controls the full scale of self-supported tumour cell motility

Our previous findings on LRP-1 and tumour cell migration and tumour formation led us to investigate the role of LRP-1 in maintaining the constitutive motility in MDA-MB-231 cells, as schematically shown in **Figure 2A**, in which we used lentiviral infection-mediated down-regulation of LRP-1. The efficiency of LRP-1 down-regulation in the cells is shown in **Figure 2B** (panel a, lane 2 vs. lane 1). When the LRP-1-downregulated tumour cells were subjected to colloidal gold migration assay, as shown in **Figure 2C**, we found that 1) the LRP-1-depleted cells have completely lost the self-supported motility under serum-free conditions (panel e vs. panel d) and 2) LRP-1 depletion did not affect the cells to respond to physiological stimulus-, such as serum-, stimulated cell motility (panel f). Quantitation of the motility data is shown in **Figure 2D**. These results demonstrate that the LRP-1 plays a commanding role in the full self-supported motility of the tumour cells.

**Figure 2.**
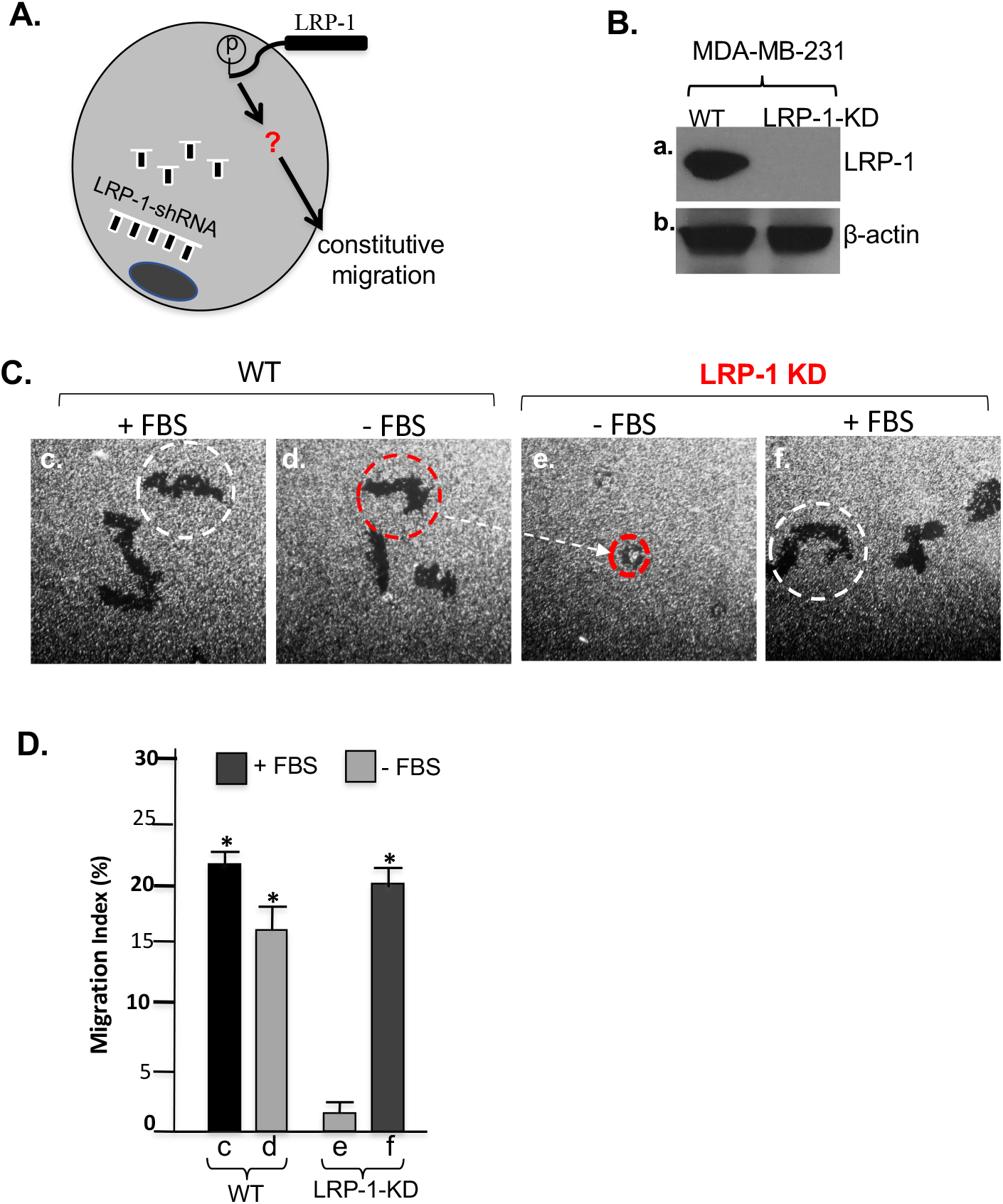
LRP-1 controls the full scale of constitutive motility of tumour cells. A) A schematic illustration of the hypothesis that cell surface LRP-1 regulates constitutive motility of tumour cells. B) Lentivirus-delivered shRNA down-regulation of LRP-1 in MDA-MB-231 cells. C) Representative images of wild type and LRP-1-downregulated cell migration under indicated conditions, in which dotter circles reflect the size of cell migrated area. D) Computer-assisted quantitation of the cell migration as Migration Index (%) (Methods). This experiment was repeated four times with reproducible results, p < 0.05.

### The “secreted Hsp90α > LRP-1 autocrine loop” only accounts for a portion of the full self-supported tumour cell motility

The commanding role for LRP-1 receptor in the self-supported motility of the tumour cells made us speculate that the previously identified “tumour-secreted Hsp90α > LRP-1 receptor” autocrine is the actual mechanism (Li et al., 2012), as schematically shown in **Figure 3A**. To study this hypothesis, we first tested whether serum-free conditioned medium of MDA-MB-231 cells stimulates motility of other cell types via secreted Hsp90. Serum-free conditioned medium was collected from either wild type or Hsp90-knockout MDA-MB-231 cells and tested for pro-motility activity using the B16 mouse melanoma cell line. As shown in **Figure 3B**, Hsp90α protein was only detected in conditioned medium from Hsp90α-wild type (wt) (lane 1), but completely absent from Hsp90α-KO (α-KO) cells (lane 2). As shown in **Figure 3C**, the conditioned medium from the wild type MDA-MB-231 cells strongly stimulated B16 cell migration (panel c vs. panels a). In contrast, the conditioned medium from the Hsp90α-knockout MDA-MB-231 cells completely lost the pro-motility activity (panel d vs. panel a), suggesting that secreted Hsp90α was a main stimulus of cell migration in the conditioned medium. Human recombinant Hsp90α protein-stimulated B16 cell migration was included as the positive control (panel b vs. panel a). Computer-assisted quantitation of the migration data is shown **Figure 3D**. However, to our surprise, neither depletion of Hsp90α inside MDA-MB-231 cells nor antibody neutralization of secreted Hsp90α function outside the cells could replicate the effect of LRP-1 downregulation: i. e. a complete blockade of the tumour cell migration. As shown in **Figure 3E**, in comparison to the effect of LRP-1 down-regulation (panel f) in comparison to the wild type cells (panel e), neither Hsp90α knockout (panel g) nor anti-Hsp90α neutralizing antibody (panel i) was able to achieve a complete inhibition of self-supported motility of the tumour cells, which was clearly indicated by the quantitation and statistical analysis of the migration tracks (**Figure 3F)**.

**Figure 3.**
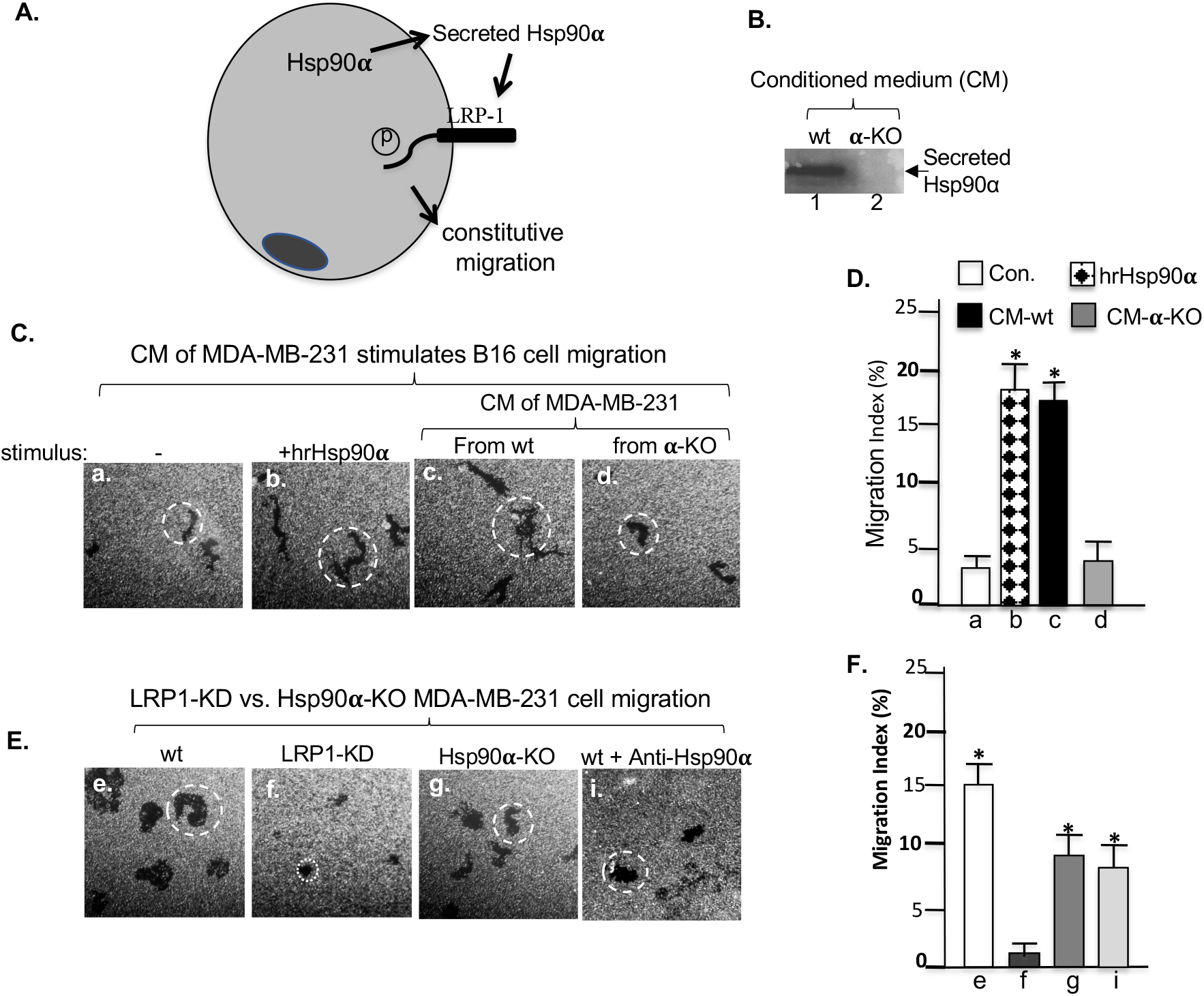
The “secreted Hsp90 > LRP-1 autocrine loop” accounts a portion of constitutive motility of tumour cells. A) A schematic illustration of the “secreted Hsp90 > LRP-1 autocrine loop” that promotes cell motility. B) Anti-Hsp90α antibody blotting of serum-free conditioned medium from wild type or Hsp90α-knockout MDA-MB-231 cells. C) Conditioned medium from wild type, bur not Hsp90α-knockout, MDA-MB-231 cells stimulates B16 melanoma cell migration, with human recombinant (hr) Hsp90α protein as control. D) Computer-assisted quantitation of the B16 cell migration. E) Hsp90α-knockout or inhibition of secreted Hsp90α by antibodies only partially reduces constitutive motility of the MDA-MB-231 cells. F) Computer-assisted quantitation of the wild type (wt) (with or without anti-Hsp90α antibody in medium), LRP-1-downregulated, and Hsp90α-knockout MDA-MB-231 cell migration. Each experiment was repeated multiple times to reach the final conclusion.

### LRP-1 stabilizes activated EGFR for the second independent pro-motility pathway

The above finding suggested that LRP-1 regulates another independent signalling pathway that, together with the “secreted Hsp90α > LRP-1 receptor” autocrine pathway, controls the full self-supported cell motility. During our antibody screening for stability of the major signalling molecules in LRP-1-downregulated MDA-MB-231 cells, we were surprised to find that EGFR (epidermal growth factor receptor) completely disappeared. As shown **Figure 4A**, the downregulation of LRP-1 (panel a, lane 2) correlated with disappearance of the ERGFR (pane b, lane 2), as well as reduced levels of phosphorylated Erk1/2 (panel c, lane 2) and cyclin D1 (panel f, lane 2). In the same cells, the phosphorylated p38 (panel d, lane 2) and Hsp90 protein (panel g, lane 2) were slightly increased, while phosphorylated Smad3 remained unchanged (panel e, lane 2). The downregulation of EGFR was unlikely due to cross reactivity of the LRP-1 shRNA with EGFR transcripts, since there was lack of any significant match to the 19-nucleotide sequence of the LRP-1 shRNA from the EGFR mRNA sequence. The EGFR in MDA-MB-231 is constitutively activated even under serum-free conditions, as indicated by anti-phosphotyrosine (PY) antibody blotting without or with EGF stimulation (**Figure 4B**, panel j, lanes 1 vs. 2). In contrast, tyrosine-phosphorylated EGFR completely disappeared in LRP-1-downregulated cells (panel j, lanes 3 and 4). Then, we tested the role of the activated EGFR in the self-supported motility of MDA-MB-231 cells by using an EGFR tyrosine kinase-specific inhibitor, Gefitinib (C_22_H_24_ClFN_4_O_3_). As shown in **Figure 4C**, even at the highest tolerable concentration of 10µg/ml prior to causing significant cell death, Gefitinib was unable to completely inhibit self-supported motility of the tumour cells, as LRP-1 downregulation does. Therefore, we postulated that only simultaneous inhibitions of both the “secreted Hsp90α > LRP-1 receptor autocrine loop” and the EGFR signalling reproduce the full effect of LRP-1 downregulation. As shown **Figure 4D**, either anti-Hsp90α neutralizing antibody (bar 5) or Gefitinib (bar 6) only partially blocked the self-supported motility (bar 3) of the tumour cells. However, the addition of both the antibody and the inhibitor showed additive effect on inhibition of the tumour cell migration (bar 7), which is insignificantly different from LRP-1 downregulated cells (bar 1). We conclude that the cell surface LRP-1 holds the pivotal position to promote the self-supported tumour cell motility by mediating both secreted Hsp90α signalling and EGFR signalling.

**Figure 4.**
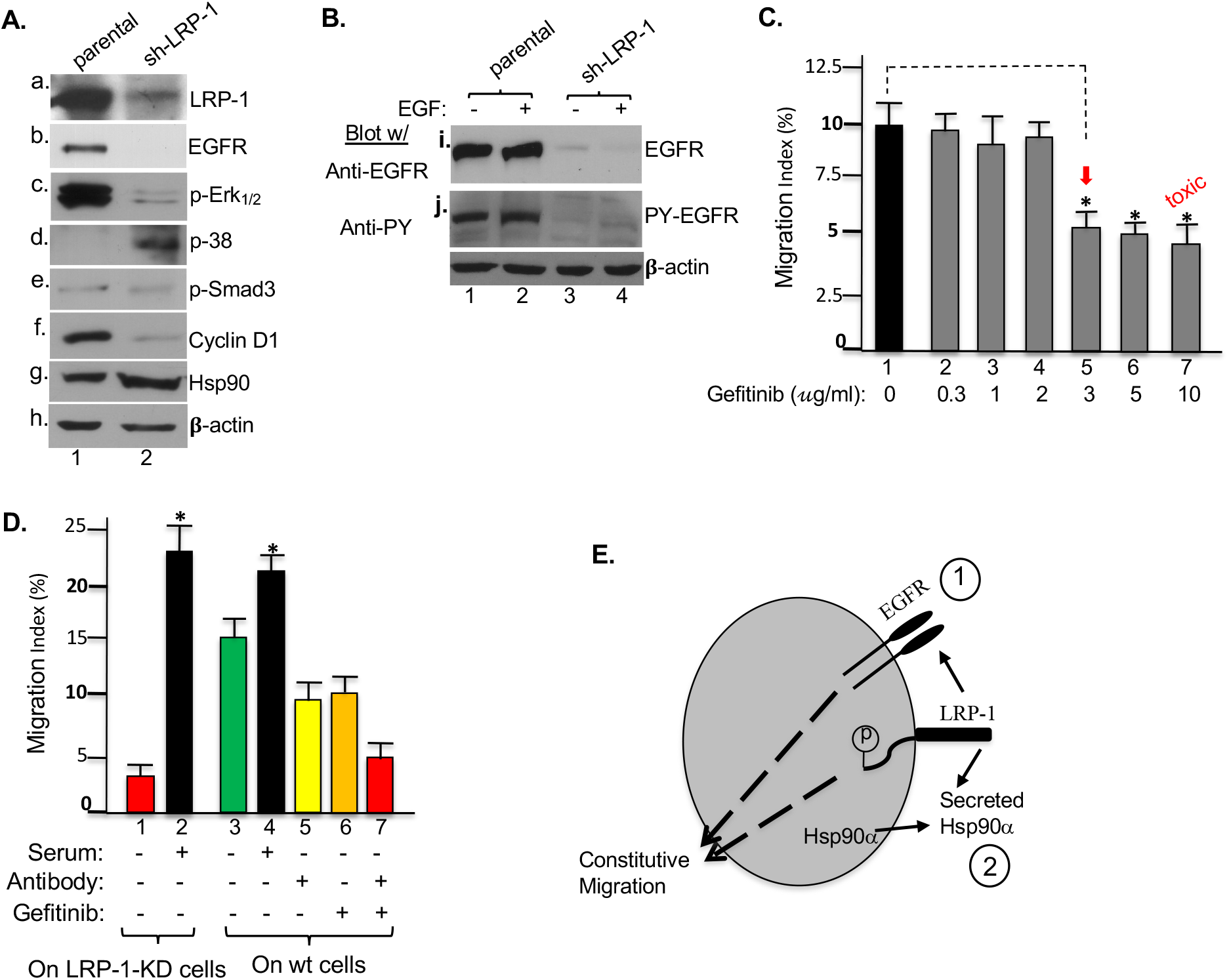
LRP-1 stabilizes activated EGFR to add the second portion of constitutive motility of tumour cells. Wild type and LRP-1-downregulated MDA-MB-231 cells were serum-starved overnight and subjected to various analyses as follows: A) Western blot shows the effect of LRP-1 down-regulation on various signalling or related proteins, as indicated; B) LRP-1 down-regulation causes disappearance of constitutively tyrosine-phosphorylated EGFR even in the absence of EGF stimulation; C) Cellular toxicity test of various dosages of Gefitinib using cell migration assay; D) Simultaneous inhibitions of the “secreted Hsp90 > LRP-1 autocrine loop” and EGFR signalling reproduces the effect of LRP-1 on constitutive motility of tumour cells; E) A working model of mechanism for constitutive motility of tumour cells. All results were reproducible in multiple repeated experiments.

## Discussion

Tumour cell motility serves as the foundation for tumour invasion and metastasis, which kill patients. Similar to the findings that different oncogenes and tumour suppressor genes drive tumorigenesis of different types of tumours, the mechanisms behind tumour cell self-supported motility likely vary among different tumours. This understanding would guide the designs of target-specific therapeutics or even personalized anti-tumour drugs. In the current study, we have taken the best-characterized human triple negative breast cancer cell lines, MDA-MB-231, as the cell model to report the first mechanism that drives tumour cell self-supported motility under serum-free conditions, a main characteristic of tumour microenvironment. In contrast to conventional wisdom of mutated oncogene-driven mechanism, our study has revealed a dual parallel pathway-participated mechanism with the cell surface LRP-1 as the pivotal coordinator. In the total absence of serum support, the “starved” tumour cells depends upon LRP-1 to turn on self-supported migration to move away from the hazard environment of no blood support. To fulfil the request, as schematically shown in **Figure 4E**, LRP-1 receptor engages tyrosine-phosphorylated EGFR signalling and, at the same time, mediates the tumour cell-secreted Hsp90α autocrine signalling, to achieve a full self-supported motility that reaches about 60% of the full cell motility under physiological conditions.

The finding that LRP-1 maintains the stability of activated EGFR cells is not cell type-specific. Similarly, another related breast cancer cell line, MDA-MB-468, has an undetectable level of LRP-1 and yet a much higher level of inactive EGFR. More convincingly, in serum-starved primary human dermal fibroblasts, EGFR remains inactivate unless EGF is added to the medium. Our experiments show that down-regulation of LRP-1 had little effect on the inactivated EGFR in the absence of EGF stimulation. However, the level of EGF-stimulated EGFR was dramatically reduced within 2 minutes in LRP-1-downregulted cells (X. Tang, C. Chang, and W. Li, unpublished data). The mechanism by which LRP-1 maintains the stability of EGFR in MDA-MB-231 cells remains to be further investigated. Since the full-length cDNA for LRP-1, which encodes for a 515-kDa extracellular α subunit and an 85-kDa trans-membrane β subunit, exceeds the maximum cloning limit for any cDNA expression vectors (Shinohara et al., 2017), it is technically not possible to do the rescue experiment with the full-length LRP-1 cDNA (Obermoeller-McCormick et al., 2001). Nonetheless, our experiments show that none of the four mini-receptor cDNAs that encodes the 85-kDa β subunit plus one of the four extracellular domains (I to IV) (Tsen et al., 2013) was able to rescue the down-regulated EGFR (negative data not shown). These data suggest that either combined domains or even the full-length wild type LRP-1 is required for stabilizing EGFR. Takayama et al reported that LRP1 controls endocytosis and c-CBL-mediated ubiquitination of the platelet-derived growth factor receptor β (PDGFRβ) (Takayama et al., 2005). It is worth investigating whether this mechanism applies to EGFR in future studies. In contrast, the second parallel “secreted Hsp90α > LRP-1” autocrine loop” to promote cell migration has been clearly established. Normal cells secrete Hsp90α only under stress, such as tissue injury signals, whereas tumour cells constitutively secrete Hsp90α driven by oncogenes, such as HIF-1α. The sole function of secreted Hsp90α is to promote cell motility during wound healing and tumour invasion. Secreted Hsp90α acts as a *bona fide* signalling protein that binds to the subdomain II in the extracellular part of LRP-1 and activates, via the NPVY motif in the cytoplasmic tail of LRP-1, the Akt and Erk1/2 kinases, to promote cell migration (Li et al., 2012). In summary, this study has revealed a novel mechanism that support tumour cells’ constitutive motility, a prerequisite for tumour invasion and metastasis.

## Materials and Methods

### Cell lines, antibodies and reagents

Limited-passage stocks of the human triple negative breast cancer cell line, MDA-MB-231, and non-transformed breast epithelial cells, HBL-100, were obtained from liquid nitrogen storage from Dr. Michael Press (University of Southern California, Los Angeles) and Hsp90α-knockout MDA-MB-231 cells were as described previously (Zou et., 2016). B16 moue melanoma cell line was purchased from ATCC. All cells were cultured in DMEM medium with high glucose supplemented with 10% FBS (Thermo Scientific, MA, USA), prior to serum-starvation and experiments. 293T cells for lentivirus production were cultured in DMEM with high glucose and 10% FBS (Thermo Scientific, MA, USA). Anti-EGFR antibody (D38B1, #4267) and anti-LRP-1 antibody (#64099) were from Cell Signalling Technology (Beverly, MA). Anti-phosphotyrosine antibody (#72) was a gift from Dr. Joseph Slessinger (Yale University). Mouse monoclonal antibodies against human Hsp90α (CA1023) purchased from Calbiochem (Billerica, MA) and Stressmarq Biosciences (Victoria, BC, Canada), respectively. The mouse monoclonal antibody, 1G6-D7, which neutralizes secreted Hsp90α function, was established in our laboratory (6). Anti-phospho-Smad3, anti-phospho-p38 kinase, anti-phospho-p44/42 MAPK (D13.14.4E, #4370) from the Cell Signalling Technology (Beverly, MA), anti-Cyclin D1 antibody (EPR2241, GTX61845, GeneTex, Irvine, CA) and anti-β-actin antibody (AC038, Transduction Laboratories, San Jose, CA) were as indicated. ECL western blotting detection reagent (# RPN2106, Amersham, Inc., Marlborough, MA). EGFR tyrosine kinase inhibitor, Getifinib (SML 1657) was purchased from Sigma-Aldrich.

### Lentivirus production and infection

The selected RNAi sequence (sense) against human LRP-1/CD91 for FG12 cloning was GACCAGTGCTCTCTGAATA. This construct was used to transfect 293T cells together with two packaging vectors, pCMVΔR8.2 and pMDG, to produce virus stocks as previously described (Cheng et al., 2008; Sahu et al., 2012).

### Western immunoblotting analysis and quantitation

Total cell lysates were separated on SDS-PAGE and transferred onto a PVDF membrane, which was subjected to Ponceau Red staining for confirming protein transfer efficiency. The primary antibodies against indicated proteins were used as instructed by corresponding manufacturers. Band intensity in Western blotting was quantified using Image J software (National Institutes of Health) via the following procedure: digital images of radiograph films were opened and converted to grayscale. Using a rectangular selection tool, a rectangle was drawn which contained all Western blot protein bands incubated with the same primary antibody from the same experiment. Next, “Plot lanes” was selected from the “Analyze” menu to create a profile plot of all the bands. Lines were then drawn between the peaks that represented darker bands. All measurements were recorded for the highlighted peaks. The peak of the control sample was selected as the standard and then the relative density of the peaks of the other bands was calculated in reference to the control. Statistical significance was determined using a two-tailed Student’s t-test and one-way ANOVA. Statistical significance was accepted when p < 0.05. Asterisk (*) was used to indicate “a significant difference vs. control”.

### Computer-assisted colloidal gold migration assay and data quantitation

Briefly, glass coverslips (35 mm in diameter) were pre-coated with 1% freshly prepared bovine serum albumin (BSA) in phosphate-buffered saline (PBS), dried by air, and placed into 12-well tissue culture plates with one coverslip per well. Colloidal gold chloride solution (colloidal gold chloride suspension (6.85 mg/ml H_2_O:30% Na_2_CO_3_ : H_2_O, 0.9 ml:3 ml : 5.5 ml) was heated in an 50-ml Erlenmeyer flask with constant swirling until boiling and then removed from the heat source. An equal volume of freshly prepared 0.1% formaldehyde solution was slowly added to the gold salt mixture with swirling. The mixture (when turning to purple brown) was immediately plated at 1 ml/well into the 12-well plates with coverslips and left undisturbed for 2 h to let the colloidal gold particles settle on the BSA-coated coverslips. The wells were gently rinsed once with 1 ml of Hanks’ balanced salt solution (HBSS) without Ca^2+^. The colloidal gold surface was coated with type I collagen in 1 ml of HBSS containing 5 mM Ca^2+^ and 20 µg/ml of native rat tail type I collagen at 37°C for 2 h. Unattached collagen molecules were removed by rinsing the wells once with HBSS without Ca^2+^. Three thousand serum-starved (0% FBS for 24 hours) cells were plated into each well and cell migration monitored following a 16 h of incubation and addition of 0.1% formaldehyde in PBS for fixation. Individual cell migration was visualized under a dark field microscope that is linked to a computer via a real-time charge-coupled device camera (KP-MIU; Hitachi Denshi, Woodberry, NY). Fifteen of randomly selected three single cell track-containing fields under each condition were photographed and quantitated by an attached computer using NIH Image 1.6 software as so-called migration index (MI). The MI represents the percentage (%) of the field area consumed by migration tracks of the migrating cells over the total field area under the microscope x 100. It is noticed the MI of the colloidal gold migration assay may vary from experiment to experiment, depending upon 1) BSA coating quality, 2) plated colloidal gold density, and 3) collagen coating and cell culture conditions. Therefore, the MIs under different conditions are only comparable within a same experiment, i.e. the actual numbers may not be comparable cross independent experiments.

### Statistical analysis

All numerical results are reported as mean and standard deviation (s.d.). Statistical significance was determined using a two-tailed Student’s t-test and one-way ANOVA. Statistical significance was accepted when *p* < 0.05. Final presentation as mean ± s.d. was based on at least three independent and corroborating experiments. Confirmation of a difference in migration as statistically significant requires rejection of the null hypothesis of no difference between mean migration indices obtained from replicate sets. The *p* value equal or less than 0.05 was considered statistically significant.

## Acknowledgments

We have neither financial nor non-financial conflict of interest. This work is supported by NIH grants GM067100 (to W. L.) and AR33625 (to M. C. and D. T. W.).

## Contribution

X. T. and C.C. participated in the design and carried out most of the experiments. D. M. helped the analyses and editing. A. S. helped om hypoxia studies. M.C. helped on primary cell culture and lentiviral vectors. W. L. designed experiments, supervised the entire project and wrote the paper with help of all above.

## Reference

1. Appert-Collin, A., Bennasroune, A., Jeannesson, P., Terryn, C., Fuhrmann, G., Morjani, H., Dedieu, S., (2017) Role of LRP-1 in cancer cell migration in 3-dimensional collagen matrix. Cell adhesion & migration. 11, 316–326.

2. Berquand, A., Meunier, M., Thevenard-Devy, J., Ivaldi, C., Campion, O., Dedieu, S., Molinari, M., Devy, J., (2019) A gentle approach to investigate the influence of lRP-1 silencing on the migratory behavior of breast cancer cells by atomic force microscopy and dynamic cell studies. Nanomedicine: Nanotechnology, Biology and Medicine 18, 359–370.

3. Cheng CF., Fan J, Fedesco M., Guan S., Li Y., Bandyopadhyay B., Bright AM., Yerushalmi D., Liang M., Chen M. et al. (2008) Transforming growth factor α (TGFα)-stimulated secretion of HSP90α: using the receptor LRP-1/CD91 to promote human skin cell migration against a TGFβ-rich environment during wound healing. Molecular and cellular biology 28, 3344–3358.

4. Chen, J.S., Hsu, Y.M., Chen, C.C., Chen, L.L., Lee, C.C., Huang, T.S., (2010) Secreted heat shock protein 90α induces colorectal cancer cell invasion through CD91/LRP-1 and NF-κB-mediated integrin αV expression. Journal of Biological Chemistry 285, 25458–25466.

5. Chen, S., Bu, G., Takei, Y., Sakamoto, K., Ikematsu, S., Muramatsu, T., Kadomatsu, K., (2007) Midkine and LDL-receptor-related protein 1 contribute to the anchorage-independent cell growth of cancer cells. Journal of cell science 120, 4009–4015.

6. Dedieu, S., Langlois, B., Devy, J., Sid, B., Henriet, P., Sartelet, H., Bellon, G., Emonard, H. and Martiny, L., (2008) LRP-1 silencing prevents malignant cell invasion despite increased pericellular proteolytic activities. Molecular and cellular biology 28, 2980–2995.

7. Herz, J., Clouthier, D.E., Hammer, R.E., (1992) LDL receptor-related protein internalizes and degrades uPA-PAI-1 complexes and is essential for embryo implantation. Cell 71, 411–421.

8. Herz, J., Strickland, D.K., (2001) LRP: a multifunctional scavenger and signaling receptor. The Journal of clinical investigation 108, 779–784.

9. Li, W., Fan, J., Chen, M., Guan, S., Sawcer, D., Bokoch, G.M., Woodley, D.T., (2004) Mechanism of human dermal fibroblast migration driven by type I collagen and platelet-derived growth factor-BB. Molecular biology of the cell 15, 294–309.

10. Li, W., Sahu, D., Tsen, F., (2012) Secreted heat shock protein-90 (Hsp90) in wound healing and cancer. Biochimica et Biophysica Acta (BBA)-Molecular Cell Research, 1823, 730–741.

11. Lillis, A.P., Mikhailenko, I., Strickland, D.K., (2005) Beyond endocytosis: LRP function in cell migration, proliferation and vascular permeability. Journal of Thrombosis and Haemostasis 3, 1884–1893.

12. Langlois, B., Perrot, G., Schneider, C., Henriet, P., Emonard, H., Martiny, L., Dedieu, S., (2010) LRP-1 promotes cancer cell invasion by supporting ERK and inhibiting JNK signalling pathways. PLoS One 5, e11584.

13. Obermoeller-McCormick, L.M., Li, Y., Osaka, H., FitzGerald, D.J., Schwartz, A.L., Bu, G., (2001) Dissection of receptor folding and ligand-binding property with functional minireceptors of LDL receptor-related protein. Journal of cell science 114, 899–908.

14. Oudin, M.J., Weaver, V.M., (2016) Physical and chemical gradients in the tumour microenvironment regulate tumor cell invasion, migration, and metastasis. Cold Spring Harbor symposia on quantitative biology. 81, 189–205.

15. Perrot, G., Langlois, B., Devy, J., Jeanne, A., Verzeaux, L., Almagro, S., Sartelet, H., Hachet, C., Schneider, C., Sick, E. et al. (2012) LRP-1–CD44, a new cell surface complex regulating tumour cell adhesion. Molecular and cellular biology 32, 3293–3307.

16. Sahu, D., Zhao, Z., Tsen, F., Cheng, C.F., Park, R., Situ, A.J., Dai, J., Eginli, A., Shams, S., Chen, M. et al. (2012) A potentially common peptide target in secreted heat shock protein-90α for hypoxia-inducible factor-1α–positive tumours. Molecular biology of the cell 23, 602–613.

17. Song, H., Bu, G., (2009). MicroRNA-205 inhibits tumour cell migration through down-regulating the expression of the LDL receptor-related protein 1. Biochemical and biophysical research communications 388, 400–405.

18. Song, H., Li, Y., Lee, J., Schwartz, A.L., Bu, G., (2009) Low-density lipoprotein receptor-related protein 1 promotes cancer cell migration and invasion by inducing the expression of matrix metalloproteinases 2 and 9. Cancer research 69,879–886.

19. Shinohara, M., Tachibana, M., Kanekiyo, T., Bu, G., (2017) Thematic Review Series: ApoE and Lipid Homeostasis in Alzheimer’s Disease: Role of LRP1 in the pathogenesis of Alzheimer’s disease: evidence from clinical and preclinical studies. Journal of Lipid Research 58, 1267.

20. Takayama, Y., May, P., Anderson, R.G., Herz, J., (2005) Low density lipoprotein receptor-related protein 1 (LRP1) controls endocytosis and c-CBL-mediated ubiquitination of the platelet-derived growth factor receptor β (PDGFRβ). Journal of Biological Chemistry 280, 18504–18510.

21. Thapa, B., Koo, B.H., Kim, Y.H., Kwon, H.J., Kim, D.S., (2014) Plasminogen activator inhibitor-1 regulates infiltration of macrophages into melanoma via phosphorylation of FAK-Tyr925. Biochemical and biophysical research communications 450, 1696–1701.

22. Tsen, F., Bhatia, A., O’Brien, K., Cheng, C.F., Chen, M., Hay, N., Stiles, B., Woodley, D.T., Li, W., (2013) Extracellular heat shock protein 90 signals through subdomain II and the NPVY motif of LRP-1 receptor to Akt1 and Akt2: a circuit essential for promoting skin cell migration in vitro and wound healing in vivo. Molecular and cellular biology 33, 4947–4959.

23. Willnow, T.E., Nykjaer, A., Herz, J., (1999) Lipoprotein receptors: new roles for ancient proteins. Nature cell biology 1, 157–162.

24. Xing, P., Liao, Z., Ren, Z., Zhao, J., Song, F., Wang, G., Chen, K., Yang, J., (2016) Roles of low-density lipoprotein receptor-related protein 1 in tumors. Chinese journal of cancer 35, 1–8.

25. Zou, M., Bhatia, A., Dong, H., Jayaprakash, P., Guo, J., Sahu, D., Hou, Y., Tsen, F., Tong, C., O’Brien, K. et al. (2017) Evolutionarily conserved dual lysine motif determines the non-chaperone function of secreted Hsp90alpha in tumour progression. Oncogene 36, 2160–2171.

